# The PRO model accounts for the anterior cingulate cortex role in risky decision-making and monitoring

**DOI:** 10.1101/2021.10.13.464327

**Authors:** Jae Hyung Woo, Habiba Azab, Andrew Jahn, Benjamin Hayden, Joshua W. Brown

## Abstract

The anterior cingulate cortex (ACC) has been implicated in a number of functions including performance monitoring and decision making involving effort. The prediction of responses and outcomes (PRO) model has provided a unified account of much human and monkey ACC data involving anatomy, neurophysiology, EEG, fMRI, and behavior. Here we explore the computational nature of ACC with the PRO model, extending it to account specifically for both human and macaque monkey decision-making under risk, including both behavioral and neural data. We show that the PRO model can account for a number of additional effects related to outcome prediction, decision-making under risk, gambling behavior, and we show that the ACC represents the variance of uncertain outcomes, suggesting a link between ACC function and mean-variance theories of decision making. The PRO model provides a unified account of a large set of data regarding the ACC.

## INTRODUCTION

The anterior cingulate cortex is a structure on the medial wall of the prefrontal cortex whose function is hotly debated. It is, nevertheless, of keen interest to psychologists because of its prominent role in processes such as reward processing, executive control and high-level thought (Heilbronner & Hayden, 2016; Paus, 2001; Ebitz & Hayden, 2016). It is also important, for the same reasons, to psychiatry, because its likely role in the progression of several diseases, including addiction, depression, and obsessive-compulsive disorder. Formalized theories and models of dACC function have been useful throughout the history of the area in driving empirical research because of their ability to generate testable and falsifiable hypotheses about functional effects.

Numerous studies have shown that the anterior cingulate cortex (ACC) and broader medial prefrontal cortex (mPFC) generates performance monitoring signals. Originally the ACC was understood to reflect error signals (Gemba et al., 1986; Niki & Watanabe, 1979), and later work suggested that ACC detected response conflict (Botvinick et al., 2001; Carter, 1998). More recently the ACC has been understood to signal the value of task options including cost and foraging value (Kolling et al., 2012; Shenhav et al., 2013; Blanchard & Hayden, 2014) and general motivation of goal-directed behavior (Parvizi et al., 2013), as well as driving loss avoidance (Boroujeni et al., 2021; Brown et al., 2007). While debate continues (Ebitz & Hayden, 2016), the ACC performance monitoring signals have increasingly been understood as reflecting prediction errors (Amador et al., 2000; Hayden et al., 2011; Kennerley et al., 2011). The PRO model (Alexander & Brown, 2011, 2014, 2019; Brown & Alexander, 2017) provides a theoretical account of how the ACC and surrounding medial prefrontal cortex (mPFC) may generate predictions and prediction errors (Figure 1A). It has been less clear how these predictions and prediction errors might drive control of behavior.

**Figure 1A.**
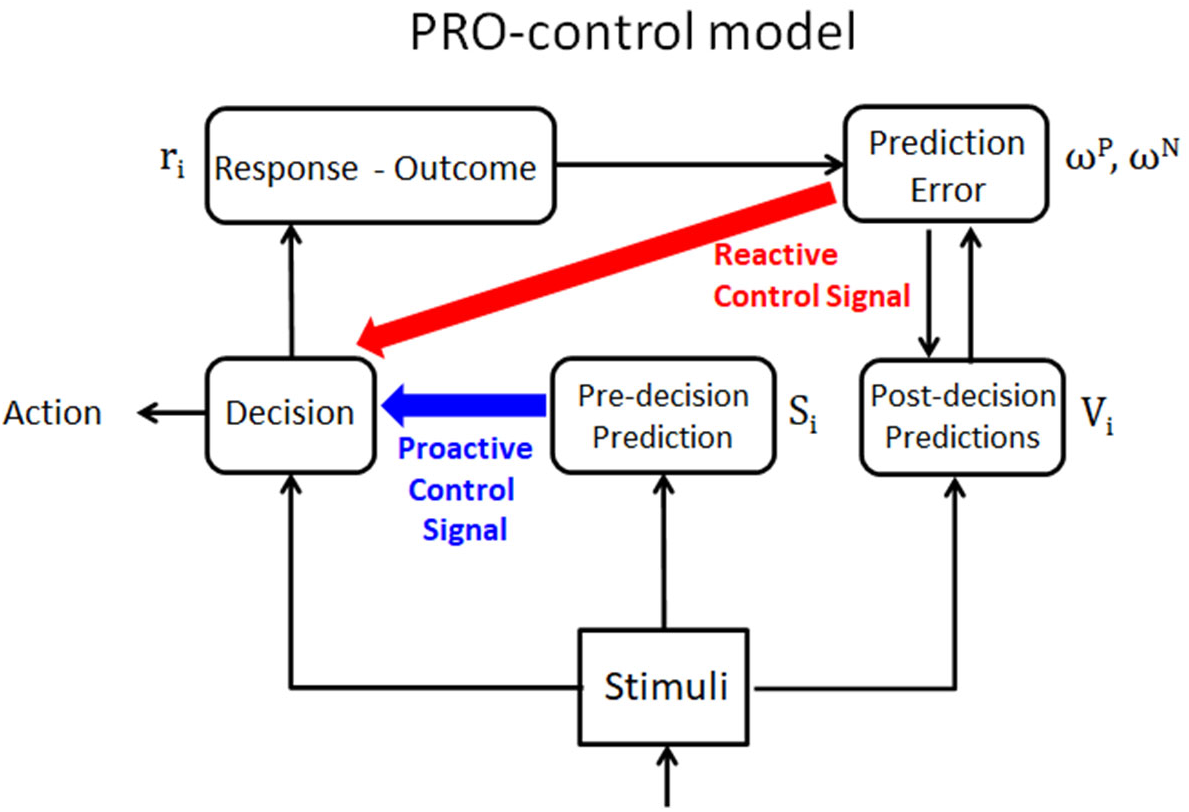
Overview of the PROcontrol model. The model (Brown & Alexander, 2017) builds on the previous PRO model by specifying two distinct processes within mPFC, namely the proactive control (blue) prior to decision and the reactive control (red) after a decision. The proactive control signal facilitates a desired response based on predecision prediction, while the reactive control signal inhibits an undesired response based on prediction error.

The PRO model, especially the PRO-control variant (Brown & Alexander, 2017), suggests two distinct control processes within the mPFC, namely a proactive control loop that controls decisions based on anticipated outcomes of responses, and a reactive control loop (Braver et al., 2012) that predicts outcomes and signals prediction errors. The proactive control processes occur *prior to* decisions, while the reactive control processes occur *after* decisions in the lead up to evaluating the consequences of a decision. Prediction errors generated by this post-decision evaluative process can lead to greater control signals in subsequent trials and modifications of the learning rate in the model.

Past simulations of the PRO-control model have shown its ability to capture numerous effects (Alexander & Brown, 2011; 2014; Brown & Alexander, 2017), but there are several key aspects of the model that have not yet been evaluated directly. One of the features of the PRO model is that it is capable of making predictions at the neuron level. That in turn means that monkey single unit data can provide important tests of PRO theory.

First, after the decision, the PRO model predicts that at the neural level, there should be a population of mPFC cells that show an upward ramping of firing rates that begin following a decision and peak at the time of an expected outcome. This corresponds to a prediction of the expected outcome. Second, before the decision, previous studies have shown that greater possible variance in the outcomes is associated with greater activity in the mPFC (Fukunaga et al., 2018), but it is unclear whether and how the PRO control model might show similar effects.

To address these issues, we perform here two distinct simulations of experiments, accompanied by novel monkey neurophysiological data. First, we developed a novel gambling task in macaque monkeys as reported previously (Azab & Hayden, 2017, 2018, and 2020). In particular, we structured the timing of trials to isolate a period during which outcome predictions are generated, by introducing a delay period (*prediction period*) after a response has been generated but before an outcome has been revealed. This affords a direct measurement of putative outcome prediction signals. Based on the PRO model, we expect to find cells with increased activity around the time a corresponding predicted outcome is likely to occur. Below we show the actual results of this task in monkeys and then explore their theoretical implications in the context of the PRO model. We find that the results are largely consistent with an account in which the ACC generates predictions and computes prediction errors, although the results necessitate some modest revision of the PRO model.

Behaviorally, we found previously that monkeys prefer options with both higher expected value and higher variance in the gamble outcomes (consistent with broader patterns seen in this species, Heilbronner & Hayden, 2013; Farashahi et al., 2019). In the PRO model, we simulate the proactive control units as responding with greater aggregate activity when the variance of an option was greater. This proactive control signal then both indicated greater variance in the gamble and biased the choice in favor of the higher variance option. This is related to mean-variance theories of decision-making (Markowitz, 1952), but with a preference to seek rather than avoid risk.

Second, to address specifically the *pre-decision* proactive control processes in humans, we revisit our previous human fMRI study with a similar gambling task (Fukunaga et al, 2018). The BOLD signal was measured while the subjects inspected the given uncertain options and made a response (*choice period*). This measurement was then regressed to three different formalisms of risk in the gamble— variance of the outcomes, probability of loss, and maximum possible loss. In PRO model, the variables corresponding to pre-decision predictions represent the pre-decisional neural activities (Figure 1A), thereby allowing a comparison with the fMRI results. Consistent with the empirical results, we find that the proactive control signals—resulting from the pre-decisional prediction activities—show significant relationship with only the variance of the gamble outcomes but not other formalisms of risk, again consistent with the monkey empirical data.

## METHODS

### i) Monkey gambling task (single-cell recording)

The task used here (Figure 1B) was used in previous studies; some of the data analyzed here have been published before; the analyses presented here are all new (Strait et al., 2016; Azab & Hayden, 2017, 2018, 2020; Maisson et al., 2021)

**Figure 1B.**
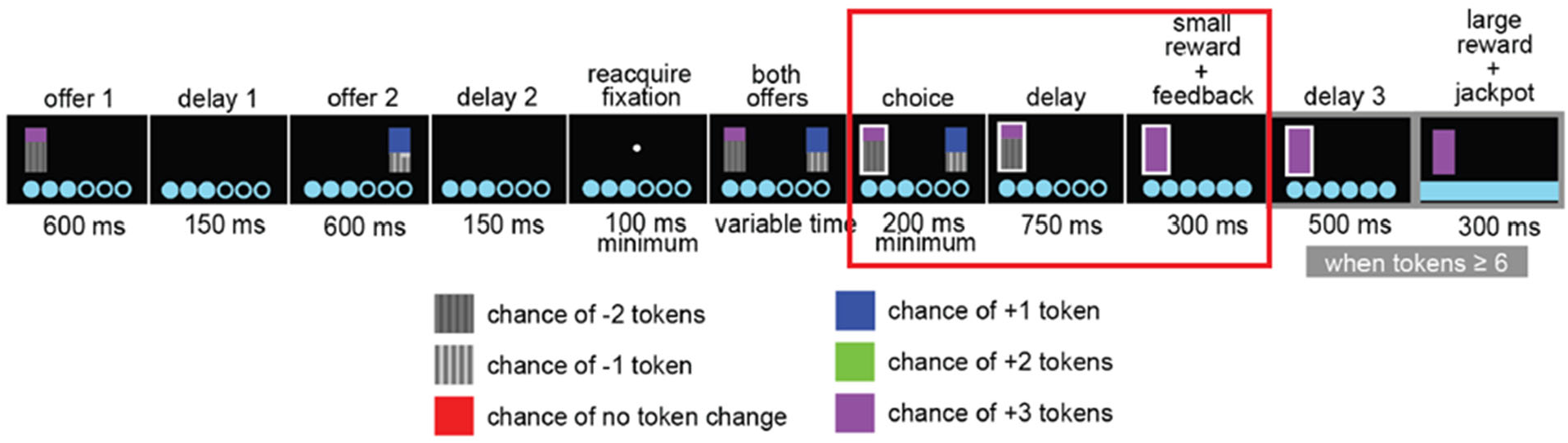
Experiment 1 task: Monkey gambling task structure to test mechanisms of post-choice outcome prediction. Monkeys were presented with each option sequentially (in a randomized order) and were made to make a choice by fixating their gaze on the desired option. The red box indicates our interval of interest: 750ms delay period right after the choice has been made, followed by 300ms of reward feedback period. (Figure adapted with permission from Azab & Hayden (2016).)

#### Surgical Procedures

All procedures were approved by the University Committee on Animal Resources at the University of Rochester and were designed and conducted in compliance with the Public Health Service’s Guide for the Care and Use of Animals. Two male rhesus macaques (*Macaca mulatta*: subject B age 5 and subject J age 6 at the start of recording) served as subjects. We used standard procedures, as described previously (Strait et al., 2014). A small prosthesis for holding the head was used. Animals were habituated to laboratory conditions and then trained to perform oculomotor tasks for liquid reward. A Cilux recording chamber (Crist Instruments, Hagerstown, Maryland, USA) was placed over the dACC and attached to the calvarium with ceramic screws. Appropriate anesthesia was used at all times; induction was performed with ketamine and isoflurane was used for maintenance. For surgical induction, we used 10–15 mg/kg of ketamine, 0.25 mg/kg of midazolam, and 2– 4 mg/kg of propofol. For maintenance, we used isoflurane, ad lib level, set depending on active monitoring procedure. For systemic antibiotics, we used cefazolin and for topical application, we used standard veterinary triple antibiotic. For analgesics, we used meloxicam, and, when judged necessary by veterinary staff, buprenorphine. Postoperative care included close monitoring and restoration of fluid intake. Animals received appropriate analgesics and antibiotics after all procedures. Position was verified by magnetic resonance imaging with the aid of a Brainsight system (Rogue Research Inc., Montreal, Quebec, Canada). Throughout both behavioral and physiological recording sessions, the chamber was kept sterile with regular antibiotic washes and sealed with sterile caps.

#### Recording Site

We approached dACC through a standard recording grid (Crist Instruments). We defined dACC according to the Paxinos atlas (Paxinos, Huang & Toga, 2000). Roughly, we recorded from a ROI lying within the coronal planes situated between 29.50 and 34.50 mm rostral to interaural plane, the horizontal planes situated between 4.12 to 7.52 mm from the brain’s dorsal surface, and the sagittal planes between 0 and 5.24 mm from medial wall (Figure 1B). Our recordings were made from a central region within this zone. We confirmed recording location before each recording session using our Brainsight system with structural magnetic resonance images taken before the experiment. Neuroimaging was performed at the Rochester Center for Brain Imaging, on a Siemens 3T MAGNETOM Trio Tim using 0.5 mm voxels. We confirmed recording locations by listening for characteristic sounds of white and gray matter during recording, which in all cases matched the loci indicated by the Brainsight system.

#### Electrophysiological Techniques

Single electrodes (Frederick Haer & Co., Bowdoin, Maine, USA; impedance range 0.8 to 4 MU) were lowered using a microdrive (NAN Instruments, Nazaret Illit, Israel) until waveforms of between one and three neuron(s) were isolated. Individual action potentials were isolated on a Plexon system (Plexon, Inc., Dallas, Texas, USA). Neurons were selected for study solely on the basis of the quality of isolation; we never pre-selected based on task-related response properties. All collected neurons for which we managed to obtain at least 250 trials were analyzed.

#### Eye Tracking and Reward Delivery

Eye position was sampled at 1,000 Hz by an infrared eye-monitoring camera system (SR Research, Ottawa, Ontario, Canada). Stimuli were controlled by a computer running Matlab (Mathworks) with Psychtoolbox (Brainard, 1997) and Eyelink Toolbox (Cornelissen, Peters & Palmer, 2002). Visual stimuli were colored rectangles on a computer monitor placed 57 cm from the animal and centered on its eyes. A standard solenoid valve controlled the duration of juice delivery. The relationship between solenoid open time and juice volume was established and confirmed before, during, and after recording.

#### Behavioral Task

Monkeys performed a two-option gambling task. The task was similar to ones we have used previously (Strait et al., 2014), with two major differences: (1) monkeys gambled for virtual tokens— rather than liquid—rewards, and thus (2) outcomes could be losses as well as wins.

Two offers were presented on each trial. Each offer was represented by a rectangle 300 pixels tall and 80 pixels wide (11.35° of visual angle tall and 4.08° of visual angle wide). 20% of options were safe (100% probability of either 0 or 1 token), while the remaining 80% were gambles. Safe offers were entirely red (0 tokens) or blue (1 token). The size of each portion indicated the probability of the respective reward. Each gamble rectangle was divided horizontally into a top and bottom portion, each colored according to the token reward offered. Gamble offers were thus defined by three parameters: two possible token outcomes, and probability of the top outcome (the probability of the bottom was strictly determined by the probability of the top). The top outcome was 10%, 30%, 50%, 70% or 90% likely on gamble offers.

Six initially unfilled circles arranged horizontally at the bottom of the screen indicated the number of tokens to be collected before the subject obtained a liquid reward. These circles were filled appropriately at the end of each trial, according to the outcome of that trial. When 6 or more tokens were collected, the tokens were covered with a solid rectangle while a liquid reward was delivered. Tokens beyond 6 did not carry over, nor could number of tokens fall below zero.

On each trial, one offer appeared on the left side of the screen and the other appeared on the right. Offers were separated from the fixation point by 550 pixels (27.53° of visual angle). The side of the first offer (left and right) was randomized by trial. Each offer appeared for 600 ms and was followed by a 150 ms blank period. Monkeys were free to fixate upon the offers when they appeared (and in our observations almost always did so). After the offers were presented separately, a central fixation spot appeared and the monkey fixated on it for 100 ms. Following this, both offers appeared simultaneously and the animal indicated its choice by shifting gaze to its preferred offer and maintaining fixation on it for 200 ms. Failure to maintain gaze for 200 ms did not lead to the end of the trial, but instead returned the monkey to a choice state; thus, monkeys were free to change their mind if they did so within 200 ms (although in our observations, they seldom did so). A successful 200 ms fixation was followed by a 750 ms delay, after which the gamble was resolved and a small reward (100 μL) was delivered—regardless of the outcome of the gamble—to sustain motivation. This small reward was delivered within a 300 ms window. If 6 tokens were collected, a further delay of 500 ms was followed by a large liquid reward (300 μL) within a 300 ms window, followed by a random inter-trial interval (ITI) between 0.5 and 1.5 s. If 6 tokens were not collected, subjects proceeded immediately to the ITI.

Each gamble included at least one positive or zero-outcome, ensuring that every gamble carried the possibility of a win (or at least no change in tokens). This decreased the number of trivial choices presented to subjects, and maintained motivation.

### ii) Human gambling task (fMRI)

In the second experiment, human participants performed a similar 2-option gambling task. The task and fMRI data have been described previously (Fukunaga et al., 2018), and we describe the task here briefly. For each trial, participants were made to choose between a *Gamble* (option 1) with three different possible outcomes, or a *SureThing* (option 2) which guaranteed a specified outcome with 100% probability. Five different gambles were constructed (Figure 1C, middle panel) in order to orthogonalize the three properties of the gamble, namely probability of loss, maximum possible loss, and variance of outcome. This orthogonalization ensures that the three formal properties are mutually uncorrelated across all gambles, so as to avoid any confounding effects for loading the fMRI activity on one of the formal property regressors. (All five gambles had the same expected value.)

**Figure 1C.**
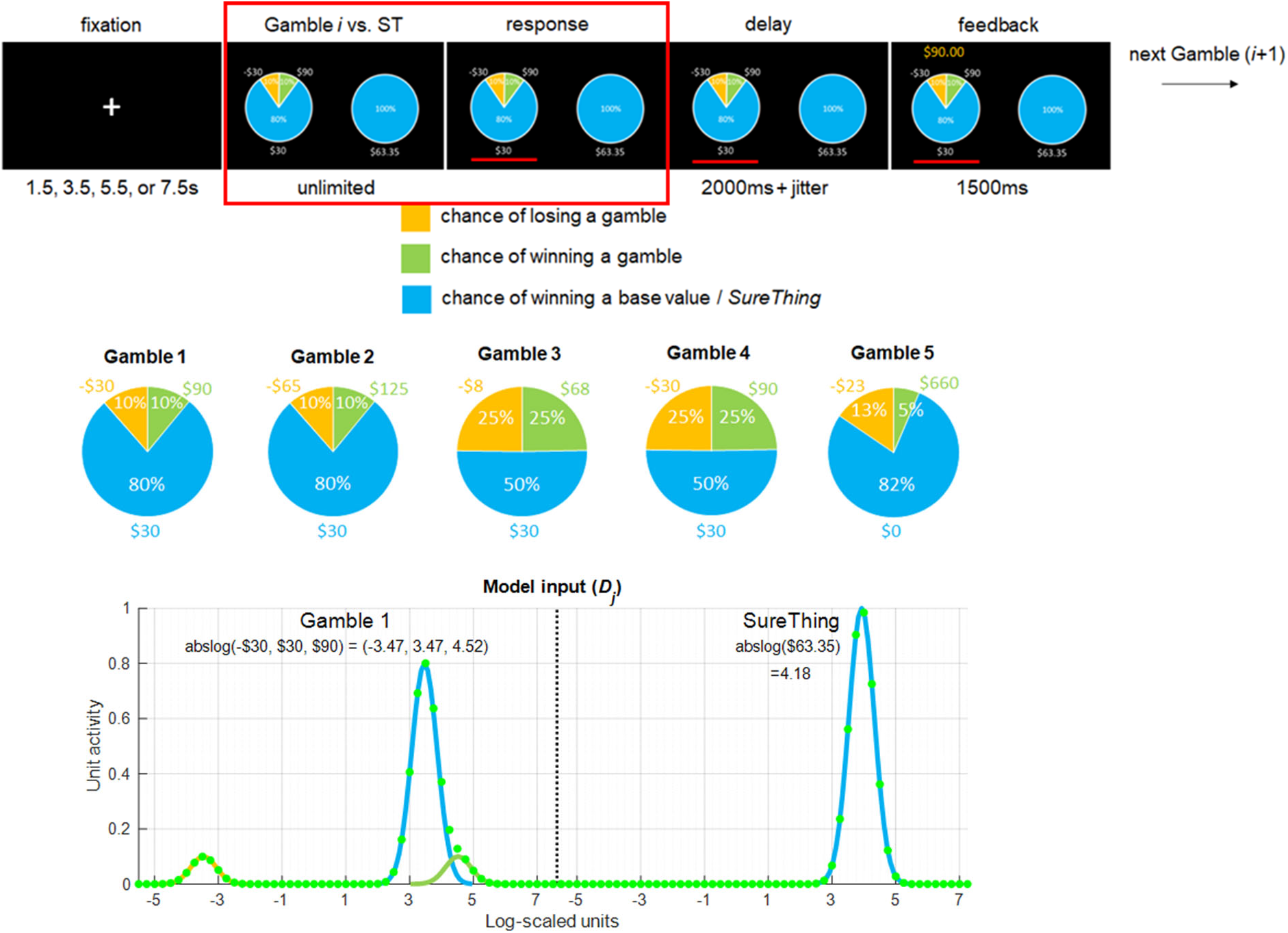
Experiment 2 task: Human gambling task design to test mechanisms of pre-choice proactive control. Each gamble is presented with its corresponding *SureThing* (ST) option specified by the linear control algorithm (see text for details). The red box indicates the pre-choice period during which a proactive control signal is generated. Middle row shows five gamble types, specifically designed to mutually decorrelate the formal properties of risk (variance of outcome, probability of loss, maximum possible loss). Each gamble is presented in each trial in a sequential order. Bottom row shows an example model input for Gamble 1 and *SureThing* options shown inside the red box. Each token value is represented by log-scaled discrete units (Nieder, 2002) (See Supplementary Materials).

The task also reduced the confounding effect of choice probability by ensuring that *Gamble* and *SureThing* options are chosen with equal probability (50%) across all trials and gamble types. This was done by using a linear control algorithm that estimated the certainty equivalent (CE) value. CE is the specified payoff amount for the *SureThing* option for which the participant equally prefers the *SureThing* and *Gamble* alternatives. The details of this algorithm can be found in Fukunaga et al, 2018. The empirical results showed that the algorithm was successful at maintaining the 50% choice probability. Furthermore, the fMRI analysis indicated that mPFC activity is significantly correlated only with the variance of the gamble offered. The regressors for other formal properties—probability of loss or maximum possible loss—were found to not load significantly on neural activity.

In light of these results, we investigate how the PRO-control model can account for the correlation between mPFC activity and the variance of the gamble outcome, and the consequent effects on cognitive control. To provide a possible mechanism for such effects, we begin with our finding below that the reaction time is the fastest for the highest-variance gamble, while the overall CE value was the lowest for the highest variance gamble. This suggests that at least in this task, a higher gamble variance is associated with both avoidance of the risky option and a faster response to the safer option. We explore this further in the computational model development below.

### iii) Computational simulations: the PRO-control model

For the simulations of the two experiments, we employ the same PRO-control model that has been reported in Brown & Alexander (2017). This model builds on the original PRO model (Alexander & Brown, 2011) and uses the same set of model parameters common to various task simulations. In essence, the PRO-control model is a generalization of standard reinforcement algorithms, and its architecture can be specified as a three-component model consisting of Actor, Controller, and the Critic. The detailed description of this model can be found in the previous work (Alexander & Brown, 2017).

In order to simulate the post-decision neural activity, we look at the Critic component of the model, which signals both unexpected events and unexpected non-events, regardless of valence (*ω^P^* and *ω^N^*, respectively). The Critic evaluates the learned predictions against the actual outcome in real time, with a process that resembles a temporal difference learning algorithm, but with some key differences as reported previously (Alexander & Brown, 2011). *ω^P^* is the positive surprise unit, which represents the prediction error due to an outcome that was not predicted to occur (surprise by occurrence); *ω^N^* is the negative surprise unit, which represents the prediction error due to a predicted outcome that failed to occur (surprise by omission). These two types of units collectively represent the prediction error signals during post-decision evaluation.

To study the effect of the pre-decision neural activities, we look at the Controller component of the model (*S*) which tracks the learned response-outcome conjunctions. In particular, we focus on the proactive control signals highlighted as the blue pathway in Figure 1A. This mechanism combines the information in the pre-decision prediction units and transforms it into weakly excitatory signals specific to each response.

A key property of the proactive control pathway is that its activity is greater when the variance of the outcomes of the gamble option is greater. The intuition behind this is twofold. First, the gamble prospects are represented by numerical receptive fields (Nieder et al., 2002). This means that a cell may respond most strongly for an anticipated outcome of e.g. winning three tokens, but it will respond weakly also for four or two tokens. Second, excitatory inputs to a given cell are underadditive here. For example, if the two possible gamble outcomes are four or five tokens, then the cell that maximally responds to five tokens will also be weakly excited by stimuli indicating the prospect of four tokens. However, and this is crucial, the cell’s firing rate will be weaker than the sum of its firing rates individually for four tokens alone or five tokens alone. This means effectively that when the variance of a gamble option is small, the corresponding outcome prediction representations are activated weakly by more than one stimulus. Thus, the summed activity across all cells will be less than the summed activity in a case when gamble outcomes have a higher variance and thus do not activate cells with overlapping receptive fields (Figure 1C). In this way, the model layer *S* shows greater summed activity over all units when the variance of a gamble prospect is higher. In turn, this signal that correlates with gamble variance can bias model decisions toward less risky options. The model is described in more detail in the Supplementary Materials.

### Experiment 1: Simulating monkey neurophysiology

#### Results

##### Monkey behavior

We examined the relationships among choice preference, expected value, and variance in the monkeys. They showed the expected preference for the higher expected value options, and they also showed a preference for the higher variance option (Figure 2). We fit the PRO model to the monkey behavioral data (Supplementary materials), and the PRO model was able to account for these effects, as shown in Figure 2. The proactive control signal was positively correlated with variance and biased the choice in favor of the higher variance options. The model simulations showed a strong positive correlation between the expected value difference and choice probability (r= 0.98, p < 0.0001), consistent with the monkey empirical data (r= 0.97, p < 0.0001), and between the variance difference and choice probability (r= 0.7169, p < 0.0001), also consistent with the monkey empirical data (r= 0.99, p < 0.0.0001) (Azab & Hayden, 2017).

**Figure 2.**
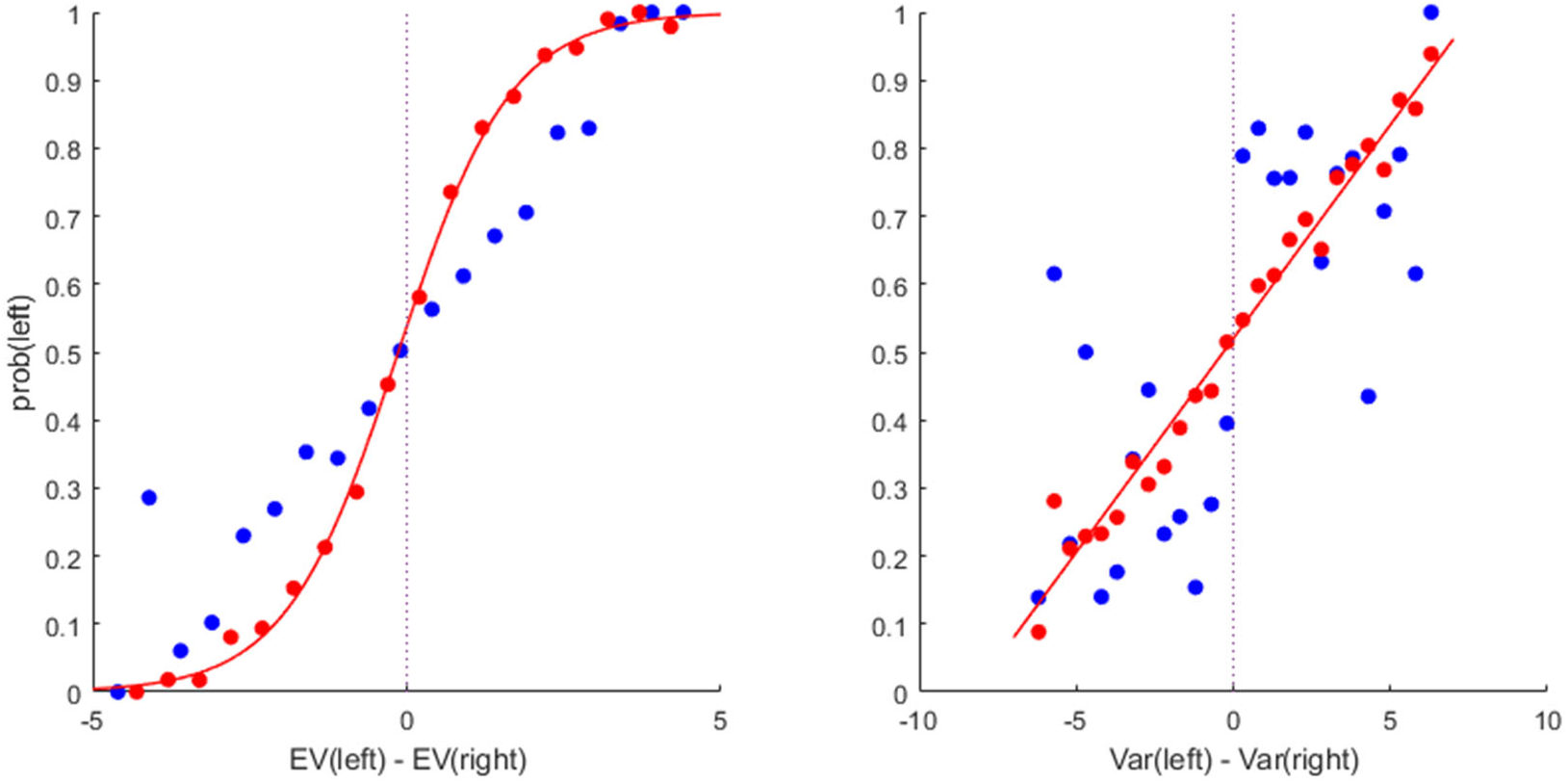
Monkey choice behavior (red) and PRO model simulation (blue). (A) Empirical choice behavior in red based on expected value difference between two options, fit to a sigmoid function. Subjects chose the left option more often as its expected value increased relative to the right option. Model simulation in blue. (B) Empirical choice behavior in red based on variance difference between two options, fit to a linear function. Subjects chose the left option more often as its variance increased relative to the right option, suggesting that the subjects were risk-seeking. Model simulation in blue.

#### Single-unit time courses

We found that single units in the macaque ACC show several distinct patterns of modulation between the decision and outcome period, as the monkey anticipates the outcomes of its response (Figure 3). Specifically, the macaque data show that a number of cells ramp *up* towards an anticipated outcome, while others show a ramp *down* towards an anticipated outcome. Our previous modeling work with the PRO model predicted consistent increasing activation towards a predicted event. This occurs as the PRO model learns a temporally structured expectation of a specific outcome (Alexander & Brown, 2011; Forster & Brown, 2011). The PRO model simulates this as the moment-by-moment temporally discounted sum of the expected future events, much as in temporal difference learning. Notably however, the original PRO model predicted only the ramp up toward the predicted event but not the ramp down.

**Figure 3.**
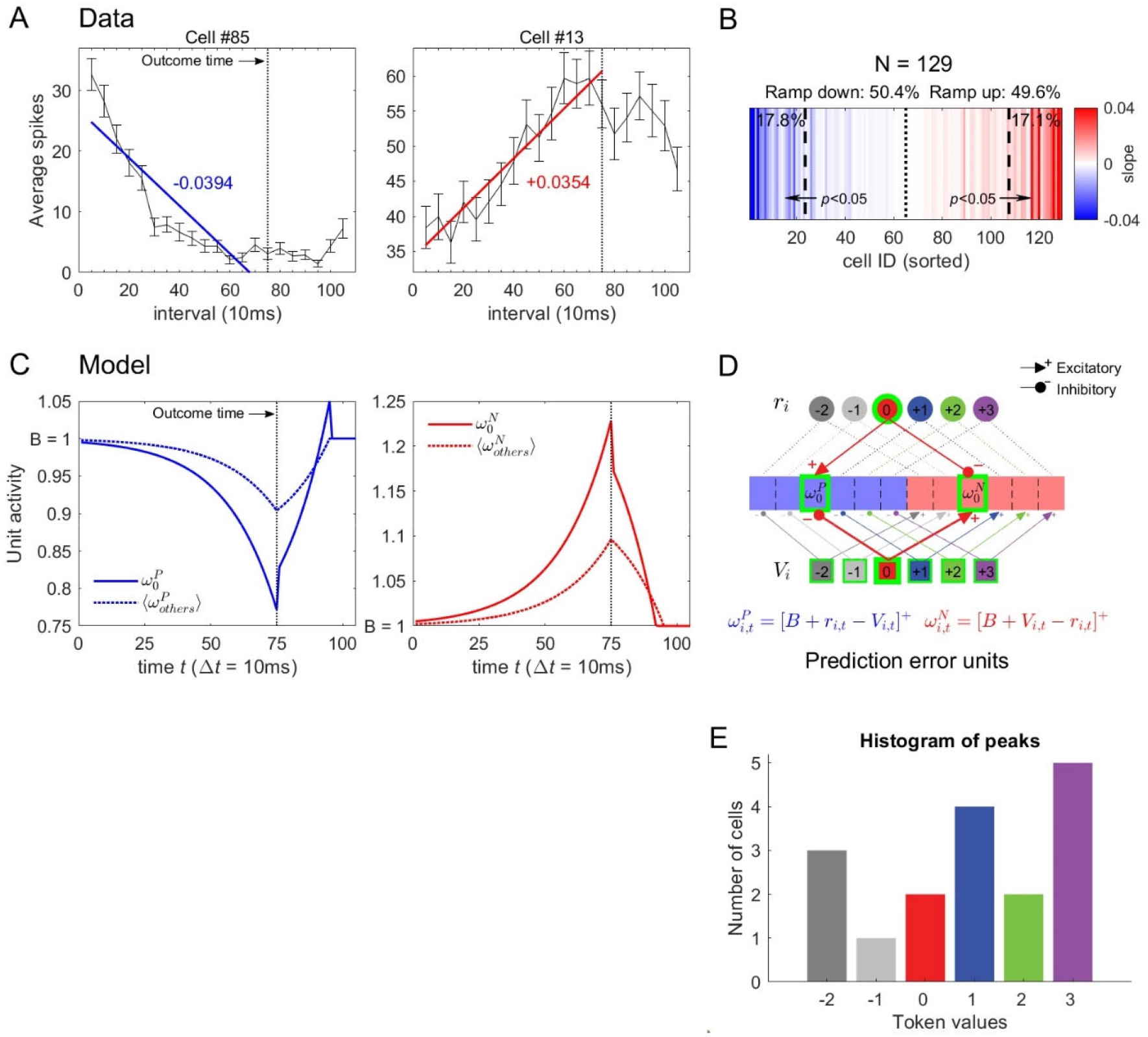
Monkey single-unit neurophysiology and the PRO model simulation of the cell behavior. (A) Example activities of dACC cells during the anticipatory period. The vertical dotted line represents the time in which the outcome is revealed. (B) The population of the cell types is represented by the heat map indicating the slope of the spiking rate (red for positive, blue for negative), showing a fairly even split between upward and downward ramping cells. (C) Example model prediction error unit activities during the anticipatory period, after which token ‘0’ is revealed. (D) Overview of the prediction error units of the PRO model. Here, the structure of the model requires the same number of positive and negative surprise units, corresponding to each possible response-outcome conjunction. (E) Histogram of the numerical receptive field peaks among monkey dACC cells, showing a range of numerical receptive fields as posited in the model.

Given the results we found in the monkey task, we explored the discrepancy between the earlier PRO-control model and the macaque results. We noted that in the PRO-control model, prediction error cells are divided into two types, namely unexpected occurrence (*ω^P^* or positive surprise) cells, and unexpected non-occurrence (*ω^N^* or negative surprise) cells. The positive surprise cells are inhibited by a predicted outcome (*V_i_*) and excited by an actual outcome (*r_i_*). Conversely, the negative surprise cells are excited by a predicted outcome and inhibited by an actual outcome. The original published PRO model made the simplifying assumption that the baseline firing rate of cells at rest is zero. In real neural cells, however, the baseline firing rate is typically small but non-zero. Given the results above, we modified the PRO-control model to simulate the more realistic non-zero baseline firing rate (*B*) at rest. This modification leads to an effect in which predictions of an imminent outcome excite negative surprise cells, causing them to ramp up above their baseline firing rate toward the predicted outcome. Crucially, predictions of an imminent outcome cause the positive surprise cells ramp down below their baseline firing rate, as the predictions (*r_i_*) inhibit the positive surprise cells (*ω^P^*). Thus, the macaque ramp-up cells and ramp-down cells above can be understood as consistent with the PRO-control model’s previously-posited theoretical positive (*ω^P^*) and negative (*ω^N^*) prediction error cells, respectively (Figure 3).

The PRO model above also posits that the monkey ACC cells will have numerical receptive fields (Nieder, 2002). To explore this, for each of the 129 cells, we created regressors representing each of the six possible outcomes. Two of these were set to 1 on a given trial, corresponding to the available outcomes, while the remaining regressors were set to zero. We then fit a general linear model to the cell’s firing rates during the prediction period (0-750ms after choice), so that the beta weights represent the firing rate of the cell when a given outcome was possible. We found 17 out of 129 cells with significant modulation as a function of available outcome, as measured by the F-statistic for the regression versus a constant model. The peak firing rates as a function of available outcomes spanned a range of values. The results shown in Figure 3E show that the monkey ACC cells indeed had a distribution of numerical receptive fields, consistent with the model assumptions. We also tried another approach in which the magnitude of the regressor for each trial was proportional to the probability of a given outcome, and the results were largely similar.

#### Neural coding of gamble properties

Next, we explored whether prediction-related cell activity after the decision was correlated with the monkey behavior, and specifically whether it correlated with certain categories of outcome, formalized as seven trial statistics in total. An important prediction of the PRO model is that the units in the model correspond to the activities of specific neurons, and therefore it is important to show that such neurons exist. These categories of outcome include: (1) Expected values of the gamble (*EV*); (2) Reward prediction error, or the difference between expected value and the outcome (*RPE*); (3) maximum obtainable token value within the given gamble bars (*maxWin*); (4) minimum winnable token value excluding the loss (*minWin*); (5) maximum amount of possible loss tokens (*possLoss*); (6) variance of the gamble (*Far*); and (7) entropy of the gamble. We compared the total predictive cell activity during the anticipatory period (defined as 350ms-750ms after a choice was made) and tested whether the activity is significantly correlated to any of the above seven formalisms. Note that Reward Prediction Error (*RPE*) is included as a control, as the cell activity is oblivious to the actual outcome which is only presented 750 ms *after* the choice. For each cell included in the dataset, we regressed the cell activity on the seven formalisms within the given trials. After obtaining the total number of significantly related cells for each formalism, we then conduct Fisher’s exact tests to determine whether the result significantly deviates from the null hypothesis that the result is due to chance.

The regression analysis showed that the macaque cell activities during the anticipatory period were significant with respect to four of the above trial statistics: expected value (*EV*), maximum winnable outcome (*maxWin*), variance (*Var*), and entropy (Figure 4A). As expected, the reward prediction error (*RPE*) showed significance for ~4.65% of the total cells (6 out of N = 129), which corresponds to the false positive rate (0.05) of the hypothesis test. We then compared the monkey empirical results with the PRO model. In Fig 4B, we regressed the averaged PRO-control model prediction unit activities (ω^N^, ω^P^) to the same seven trial statistics from the simulated gamble. Note that the overall proportions of the significantly correlated units are higher in the model simulation because there is no between-subject variance in the model unit cell activities, which would otherwise widen the distribution around the population means. Similar to the empirical data, the model prediction unit activities were most strongly associated with expected value (*EV*), maximum winnable outcome (*maxWin*), and variance (*Var*) of the simulated gamble. The larger proportion of cells that show a positive correlation between gamble variance in particular and neural activity is consistent with the PRO model results predicting greater aggregate ACC activation with greater gamble variance.

**Figures 4.**
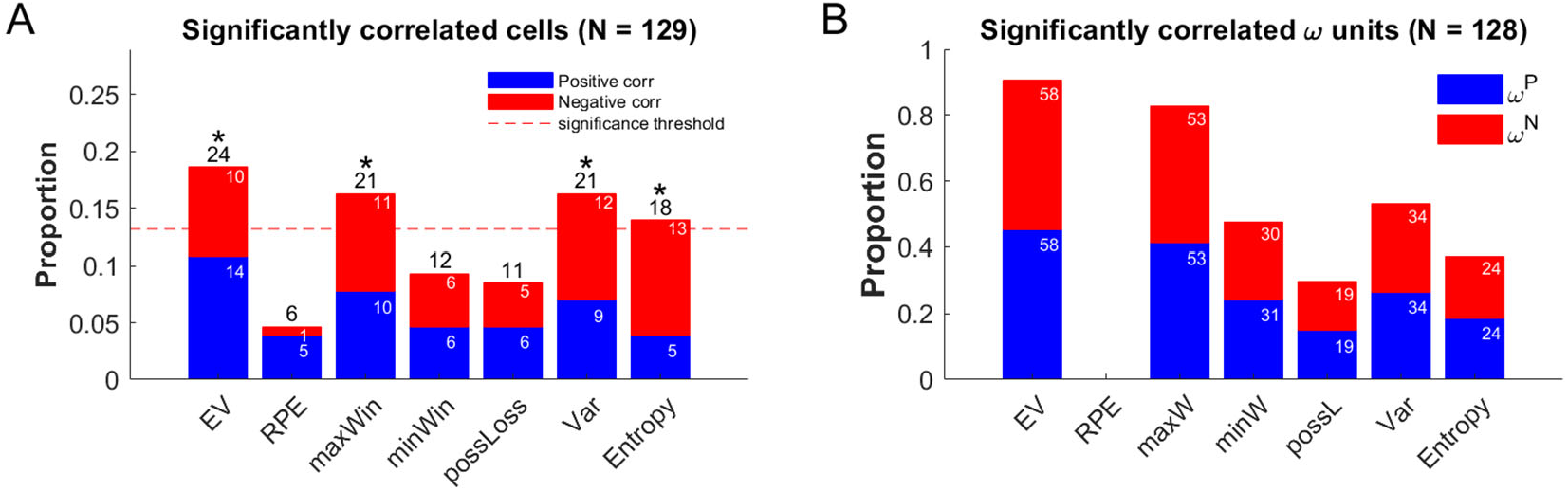
Percentage of significantly correlated cells to each trial statistics. (A) Monkey neurophysiology. The red dotted line indicates the threshold for significance correlation in the Fisher’s exact test. Four trial statistics—EV, maxWin, Variance, and Entropy—were shown to be significantly correlated with prediction cell activity (*p* = 0.05). Blue stack indicates positively correlated cells, red stack negatively correlated. (B) Results from the model prediction error units (*ω^P^* and *ω^N^*). The units show approximately the same pattern seen in the monkey data, with the strongest correlations among EV, maxWin, and Variance.

### Interim discussion

Previous studies on ACC activity have been primarily focused on the time period during choice and after the outcome (Hayden et al., 2011; Kennerley et al., 2011; Matsumoto et al., 2007). Yet, the anticipatory activity in ACC has been less understood (but see Blanchard et al., 2015). Here we focused on the pre-outcome activity in the monkey dACC cells and showed that individual cells can encode for higher-order statistics of the environment, as shown by the regression analysis. These results suggest that monkeys generate predictions at the neural level related to likely task outcomes. Furthermore, the PROcontrol model provides a theoretical account for how a mechanism that detects both the unexpected presence and absence of outcomes can explain the anticipatory activity in the monkey dACC cells.

It must be noted, however, that in the macaque cells we did not find sufficient evidence that the individual neuronal level response showed effects of the gamble outcomes, whether better or worse than expected, after the outcome was presented. Therefore, we tentatively conclude that as predicted by the PRO-control model, prediction and prediction error signals may be localized at different regions within dACC (Jahn et al., 2016). We hypothesize that the prediction activity (*B – V_i_, B + V_i_*) (Supplementary Materials) is localized more ventrally within dACC, while integration with the current outcome to encode prediction error (*B – V_i_ + r_i_, B + V_i_ – r_i_*) is localized more dorsally. This hypothesis is in line with the robust dissociation from a neuroimaging study (Jahn et al. 2016), where dorsal region was associated more with prediction error, whereas ventral regions was associated more with prediction of pain and value. It is possible that by recording from more ventral regions of dACC, we might not see prediction error signals. Overall, our results are consistent with specific predictions of expected value and maximum possible win amount, but also with predictions of other decision parameters related to risk, including variance and entropy, which might better guide decisions. Additionally, our computational simulations capture these effects and account for them as predictions about the value of various decision options as a basis for cognitive control over decision-making.

It remains to be explored how these anticipation-related signals might be instantiated and in turn contribute to control in humans. To explore this, below we consider a gambling task in humans, who on average tend to avoid higher variance gambles rather than seek them as macaques do (Figure 2). In the course of this, we will explore the nature of control, and whether the control signals on behavior are dominated by inhibition or excitation. Somewhat surprisingly, we find below that in the model, excitation dominates, in agreement with Parvizi et al. (2013) and Bliss-Moreau et al. (2021).

### Experiment 2: Simulating human fMRI experiment

#### Results

The recent study by Fukunaga et al. (2018) showed that greater possible variance in the outcomes of a gamble choice is associated with greater activity in the ACC and the surrounding mPFC during the period while the decision is being made (Figure 6A). That article proposed that prediction-related ACC cells may have a kind of receptive field (Nieder et al., 2002), such that a wider range of possible outcomes would lead to activation of more distinct populations of prediction cells. In effect, this would lead to a greater overall summed activation and thus explain the origin of the variance signal, consistent with the model mechanisms of variance signaling posited above in the description of Experiment 1. Experiment 2, a simulation exercise, begins with this proposal and explores whether and how it might account for both neural activity and behavior. The behavioral data from humans are shown in Figure 5A (Fukunaga et al., 2018), and the results of the simulation are shown in Figure 5B. The model was fitted behaviorally to the average Certainty Equivalence (CE) values of the five gambles from the empirical data (Fig 5A), as described in the Supplementary Material. The model CE was calculated as the mean *SureThing* (ST) value to which the model converged in the last 50 trials, out of total 300 simulation trials with 25 different random seeds (see Supplementary Methods for more details). The comparison of the Reaction Time (RT) results between the model vs. human behavioral data shows that, although the PRO-control model was fitted to the Certainty Equivalence (CE) values only, the model is able to capture the reduced RT for the fifth gamble which has a significantly large variance compared to the other four gambles.

**Figure 5.**
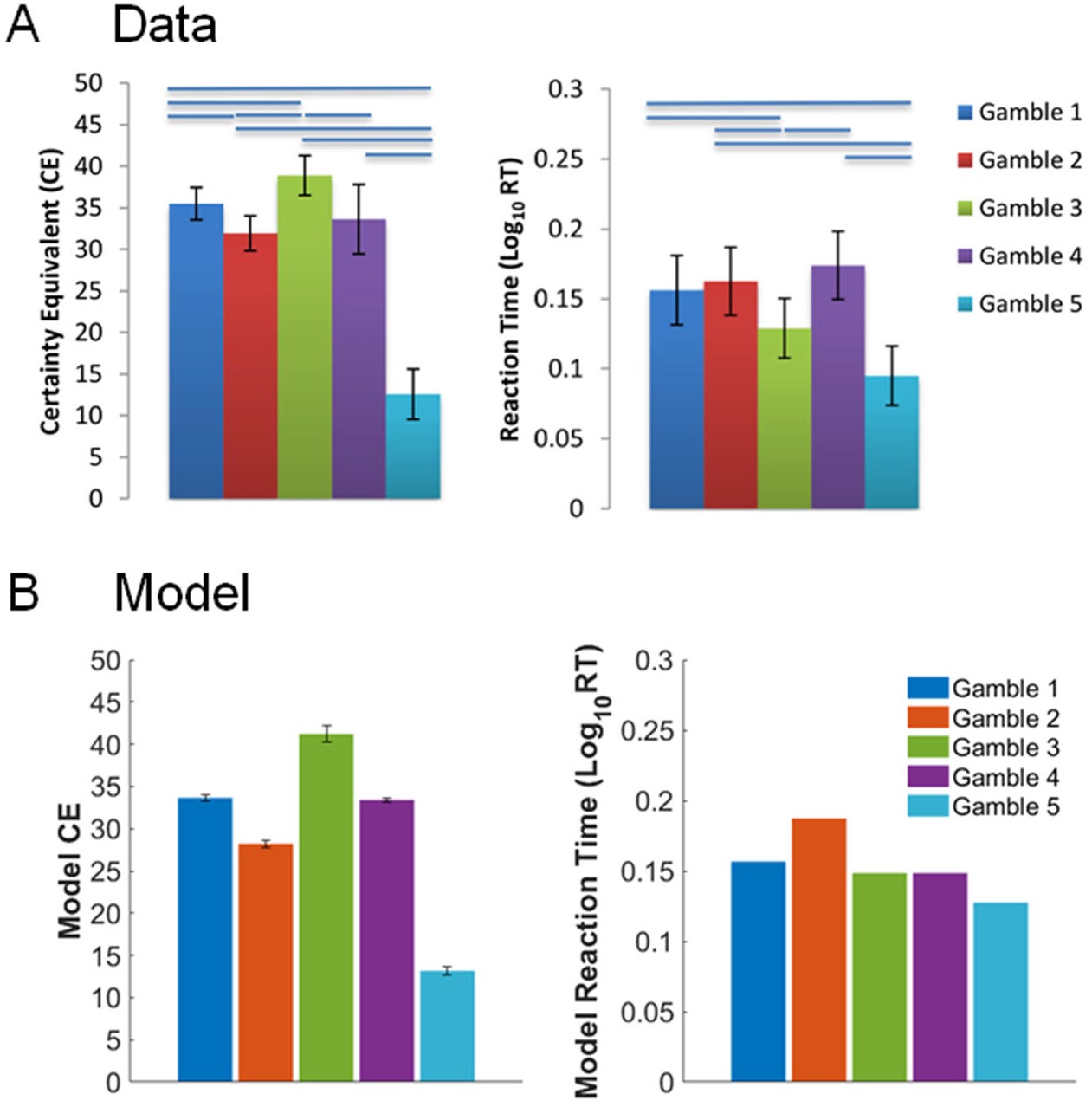
Empirical and model Certainty Equivalence (CE) and reaction times (RT). (A) Human behavioral data (N = 25). Left: Average Certainty Equivalence (CE) values for each gamble. Right: Average Reaction Times (RT) for each gamble. The fifth gamble, which has the largest variance, had both the lowest CE and RT. (B) Model results. Parameters were fitted to the average CE values only. Panel A adapted with permission from Fukunaga et al. (2018). Horizontal bars above panel A bars represent significant pairwise differences.

As with the monkey data above, the PRO model is able to account for the correlations between the variance of a gamble option and both behavioral and BOLD effects in humans. Figure 6 shows the comparison of the human BOLD data with the simulation results. The previous fMRI analyses showed that the Variance regressor showed increasing loading in the right Medial Frontal Gyrus including the ACC (Fig. 6A, red) and right Middle Frontal Gyrus/IFG (Fig. 6A, green). We simulated the proactive control-related activity as the sum of activity during the decision period in model neurons of the (S^P^) layer. Consistent with the empirical BOLD result, in our PRO-control model the summed proactive control signal (S^P^) showed the largest positive loading on the Variance regressor (p < 0.001), while showing reduced effects from probability of loss (pLoss) and maximum possible loss (MaxLoss). As with behavioral results, the BOLD simulation results are from an average from 25 different random seeds of the simulation. Note that the minimal error bars for the model (Figure 6B) are due to the lack of intersubject variability, as the model parameters were fitted to the average CE values.

**Figure 6.**
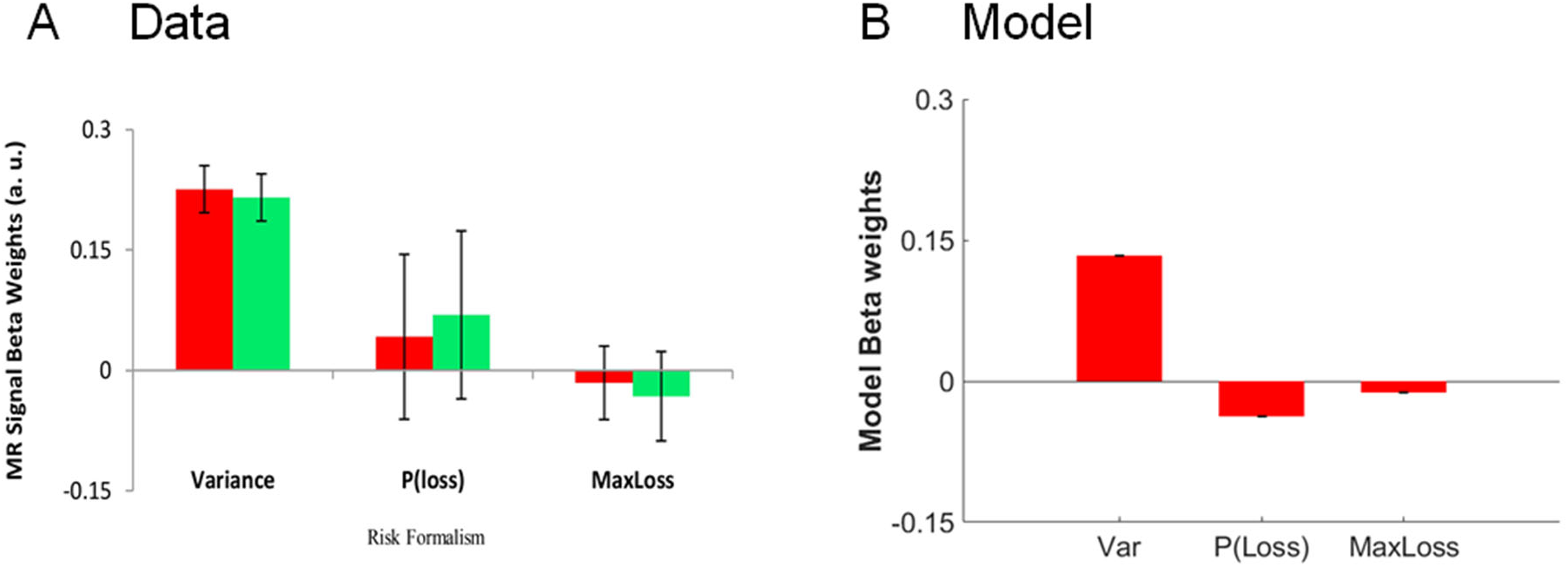
Significant loading during choice period. Risk formalism regressor during choice. (A) Human neural data. Top left (red) indicates ACC region defined by significant loading on Variance regressor (Var). Top right (green) indicates IFG/AI also loading on the variance regressor. Other regressors were not significant. (B) Simulation results also showed significant loading to variance only. Panel A adapted with permission from Fukunaga et al (2018).

## Discussion of Experiment 2

In this modeling study, we aimed to simulate both the behavioral data and neural correlates of risk, formalized as the variance of the possible outcomes. The previous study by Fukunaga et al. (2018) showed that the variance was a primary representation of risk, more so than loss probability or maximum possible loss, at least in the insula/IFG and mPFC region including the ACC. Here we show how these effects might come to be. Specifically, the sub-additive interaction of neural activity representing likely but uncertain outcomes provides a computational mechanism for the source of the variance signal, such that when the receptive fields overlap, there is less overall summed activity. In effect, the model speeds up responses when variance is higher (lower RT), while also reducing risk appetite (lower CE).

Furthermore, our results shed light on a mechanism by which the proactive control effect could operate anatomically in the mPFC region. The proactive control effect could either be excitatory or inhibitory in the PRO-control model, yet here we show that the best fit to the data involves more *excitatory* control signals, biasing decisions toward a less risky option. If control is driven from the mPFC to cause effects in the DLPFC, our results suggest that the effect of control is at least as much activation as inhibition. This appears somewhat at odds with the findings of Medalla & Barbas (2009), who reported that medial to lateral PFC projections are primarily inhibitory in rhesus monkeys, but it is consistent with the medial-lateral functional connectivity observed in a hierarchical task in humans (Zarr & Brown, 2016). The greater role of the excitatory signals as predicted by the model is consistent with the view that ACC provides a motivational signal toward a goal (Holroyd & Yeung, 2012; Kolling et al., 2012; Parvizi et al., 2013; Walton et al., 2003). Our modeling results suggest a resolution of how to reconcile motivation of extended behavior with inhibitory control: mPFC activity that drives risk avoidance does so not by inhibiting the risky choice but rather by exciting representations of the alternative, safer choice.

It is also worth noting that in the PRO-control model, the prediction and prediction error units are separate from the control pathways. This is consistent with the dorsal/ventral distinction such that prediction errors are found more dorsally in BA32/8 while affective responses are found more ventrally in BA24 (Jahn et al., 2016). This distinction is also suggested by Kolling et al. (2016) and described in Brown & Alexander (2017).

## Conclusion

In order to elucidate possible mechanisms of decision-making involving risk, our study has investigated two separate processes: *post-decision* evaluative process and*pre-decision* control process. The monkey neurophysiological data and the model simulation during the post-decision period suggest that predictions about the value of various decision options are encoded at the neural level. Furthermore, the results during the pre-decision control period clarify how ACC influences risk appetite and RT through both excitatory and inhibitory signals, with an emphasis on the former. As a whole, the PRO control model provides a broad account of ACC function and existing empirical data, as shown both above and in previous work (Alexander & Brown, 2011; 2014; Brown & Alexander 2017).

## Author contributions

BH and JWB designed the task. BH, AJ, and HA collected the macaque data. HA, BH, and JHW analyzed the macaque data. JHW analyzed the human data. JHW and JWB designed the model. JWH simulated and fit the model. JHW, AJ, and JWB wrote the paper. All authors edited and approved the final version.

## Acknowledgments

We would like to thank Meghan Castagno, Marc Mancarella, and Caleb Strait for assistance with data collection. We thank Priya Modak for helpful comments. This research was funded by a National Science Foundation grant (grant number: BCS1253576) received by BYH, and a National Institutes of Health grant (grant number: DA038615) received by BYH.

## Supplementary Materials

### Model methods

#### 1. Description of equations and PRO model modifications from Brown & Alexander (2017)

The original PRO model accounted for a number of data points across species and modalities (Alexander & Brown, 2011, 2014), and we modified it as the PRO-control model in a subsequent paper to further account for control effects (Brown & Alexander, 2017). Here, we further extend the model with some small modifications to address the two different human and monkey data sets that are the subject of PRO model simulations in the present paper. Except as noted, these changes were applied to the model to simulate both the monkey and the human data.

##### Baseline level of excitation

We changed the model to simulate the monkey data as follows, by changing Equations (15) and (16) of Alexander and Brown (2011) to:

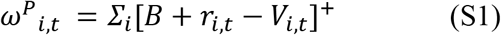

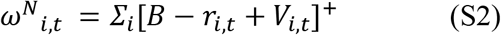

The change provides a baseline level of excitation *B*, so that inhibition would cause a substantial reduction in activity, in order to simulate the ramp down effect (Figure 2). With the previous model version, there was no baseline activity, so it was not possible for the model unit activity to decrease below zero.

##### Receptive field-like representation of gamble values

For the simulation of gambling tasks, we make the following adaptations to the PRO-control model to perform the task. First, for the stimulus representation, we use numerical receptive fields (Nieder, 2002) that are selectively activated by the incoming stimulus values.

For simulating the monkey data, the number values were represented on a linear rather than log scale. We first generate a Gaussian distribution with a mean centered on the obtained token values, where the peak of the distribution corresponds to the gamble probability. Then we select seven discretize bins for each gamble token, where the two bins at the ends are shared with the adjacent tokens to allow for overlapping activation between adjacent token values. Each of the total 32 units has activation values between 0 and 1. For the human gambling task only, since the possible gamble outcomes range across a large span of numerical values (−65 to +660), we first map the units on a log-transformed scale and then for the receptive field, we generate a Gaussian distribution with a mean centered on the obtained token values, where the peak of the distribution corresponds to the gamble probability. More specifically, each obtainable token value *X* is mapped onto the log-scaled units with a mean *m* centered at

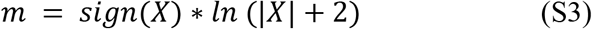

and standard deviation of 0.4. An arbitrary constant of 2 is added to the |*X*| above to prevent a logarithm of zero. The activation at unit bin corresponding to *m* is proportional to the probability of the gamble, such that the height of the Gaussian distribution in the receptive field equals the gamble probability (or 1.0 for *SureThing*). This produces the receptive field-like representations of obtainable outcomes as represented in Figure 1C (bottom panel).

##### Explicitly instructed stimulus values

Next, we assume that the subjects were explicitly aware of the probability of each gamble outcome. This is so because monkeys were extensively trained to associate the area of the gamble bar with its respective outcome probability, thus the visual information having a direct link to the gamble probability without the need to learn the association throughout the experiment. For human fMRI study, the participants were directly instructed that the probabilities of outcomes are specified by the percentages in the gamble options (Figure 1C). This modification is implemented in the model by changing Equation (1) of Brown and Alexander (2017) to set the prediction weight matrix as static and identical to the presented stimulus:

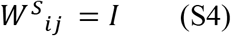

In effect, stimulus prediction weights *W^S^* do not need to be learned through experience. The weights are simply the identity matrix *I*, so that *S_i_* directly matches the presented stimulus value *D_j_*.

##### Sub-additive signal function

To account for the possible mechanism of the variance signal, we introduce a sub-additive filter to the receptive fields representing the incoming gamble stimulus values. This produces the desired sub-additive interaction of neural activity representing likely but uncertain outcomes, such that when more distinct populations of stimulus representation units are activated, there is a greater overall summed activity. This is achieved by introducing a signal function *sigact* which sensitizes the model to the low-valued input below 0.43, as described by the equation S5 below:

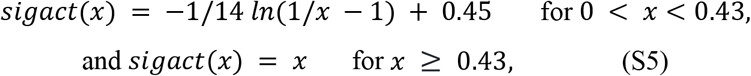

where *x* is the net input value from each unit bin. Both monkey and human gamble simulations used the same signal function. The function is plotted in Figure S1.

**Figure S1.**
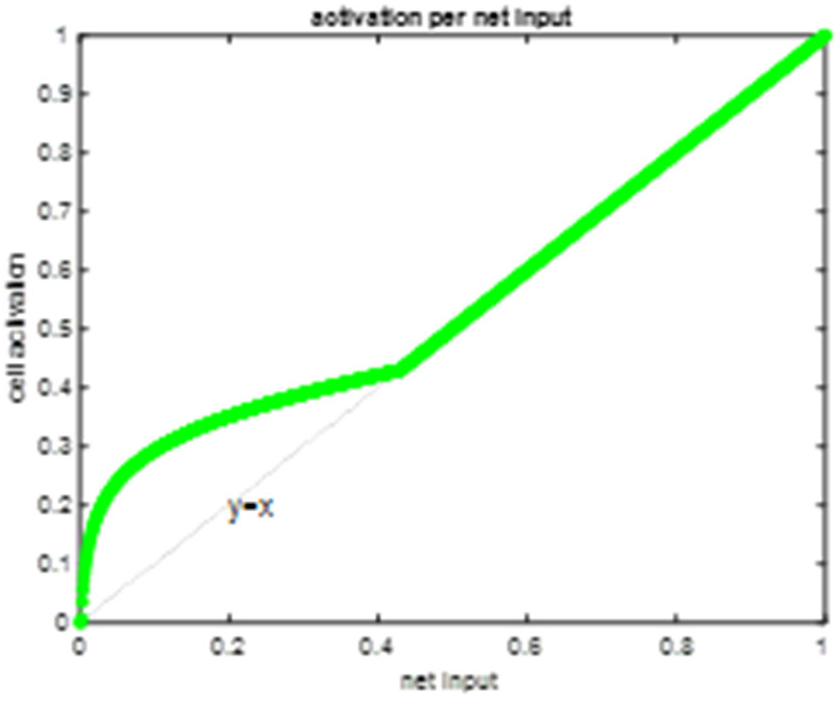
Sub-additive signal function that underlies the positive correlation of variance and summed neural activity.

##### Proactive control signal

The prediction of response-outcome conjunctions (*S_i,t_*) is filtered through this signal function and then summed to be used as the scalar source of the proactive control signal, defined as

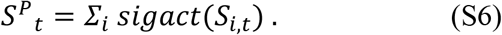

The Equations 2 and 3 of Brown and Alexander (2017) represent excitatory and inhibitory input to the response controller *C_n_*. These inputs were modified as:

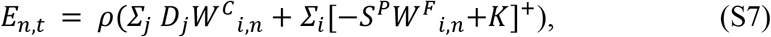

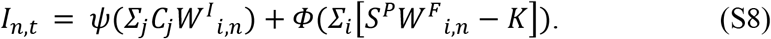

The purpose of this change was to add a threshold term *K*, in order to allow a better fit of how the proactive control signal influences behavior. Note that we omitted the reactive term *W^ω^* for these simulations as we were simulating the proactive control effects prior to outcome feedback, so reactive control does not come into play here and would have no effect. *W^C^* are prespecified weights describing the default hardwired responses by task stimuli, in the absence of the proactive control signal. For monkey gambling task simulation these weights are described as

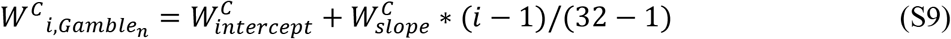

where *i* indexes each of the total 32 unit in the receptive fields representing the incoming stimulus values, so that higher token values lead to higher probability of choosing the gamble option which contain such tokens. *n* indexes the gamble option 1 and 2. The values for slope and intercept resulted from the model fitting process and are provided in Table S1. For human gambling task simulation these weights are described as

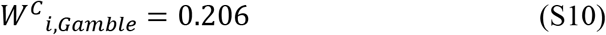

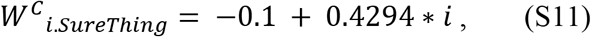

where *i* indexes each unit in the receptive fields representing the incoming stimulus values, such that greater monetary prospects are associated with larger values of *i*. Equations (S10) and (S11) resulted from the model fitting process and show that in this case, prospective gamble wins lead to a probability of choosing the gamble, but the probability is insensitive to the magnitude of the prospective win. The probability of choosing the *SureThing* option increases with increasing value of the *SureThing* prospect.

The learning rule for adjustable proactive control weights *W^F^* is modified from Equation (14) of Alexander & Brown (2011) to the following. For monkey gambling task simulations, *W^F^* was simply a fixed set of weights specifying whether the representation of one gamble provides excitatory or inhibitory input to the control unit for the respective option or the alternative. The two values resulted from the model fitting process and are provided in Table S1. For human gambling task simulations,

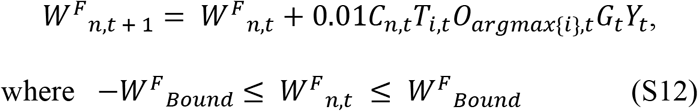

where *W^F^_Bound_* is a constant parameter that bounds the *W^F^* weights to a certain magnitude, which prevents the weights from accumulating the values indefinitely over trials. Note *O_argmax{i}_* was changed from *O_i_* from the previous work to adjust for the log-scaled transformed stimulus bins, based on the one-hot representation of discrete outcome values. Here, *O_argmax{i}_* simply represents the outcome value that was actually received. The valence *Y*, which reflects the affective evaluation of the observed outcome, is proportional to the magnitude of the token using the following equation:

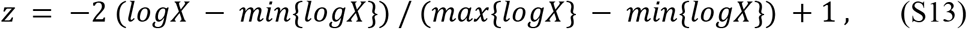

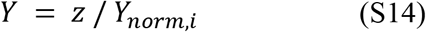

where *X* is the value of the obtained token in the gamble, and *z* is the normalized value falling in the range of [-1, 1]. *Y_norm,i_* is a constant parameter that further normalizes *z*-value according to the outcome of the gamble (*i*), defined for each scenario of obtaining a win, i.e. outcome greater than 0 (*Y_norm, i_*=1= 100) or loss, i.e. outcome less than 0 (*Y_norm, i_*=2 = 36). These values were fit as free parameters showing that for the learning laws, wins led to stronger weight updates than losses.

#### 2. Description of fitting data to the model

##### Monkey gambling task

For the monkey data, we first fit a logistic regression function between the EV difference and the probability of choosing the left option from all the monkeys’ choice behavior, and a linear function between Variance difference and the probability of choosing the left option. We then fit the PRO model parameters to the regression functions derived from the original monkey data, with the model configured to accept task-specific input as shown in Figure S2.

##### Human risk types task

The human task is described in Fukunaga et al. (2018). In the task, the staircase algorithm for determining *SureThing* (ST) values was implemented as follows: If the model or subject chose the Gamble option for a given gamble, the ST value was increased by $1.5 on the next trial with that gamble, and if the subject chose the ST option, then the ST option was decreased by $1.5 on the next trial with that gamble. All initial ST values for all gambles were $30 to match the expected value of the gamble.

For calculating the Certainty Equivalence (CE) from the model simulations, we ran a total of 300 gamble trials and took the average *SureThing* value of each gamble from the last 20 trials as the model CE.

All new parameters for this task simulation were fit by a grid search, and the results are shown in Table S1 below.

#### 3. Table of parameters used

The model was fit with a grid search over the parameter space separately for monkey and human data, to capture both neurophysiological effects and behavioral data. The results of the parameter fits are below, for monkeys (Table S1) and humans (Table S2).

**Table S1.** Model Parameters (Monkey) A&B: Alexander & Brown (2011) for the original PRO model. B&A: Brown & Alexander (2017) for the extended PRO-control model. All model parameters that appear in the previous studies are kept the same and are denoted by an asterisk (*).

| Parameter | Description | Value | Equation |
| --- | --- | --- | --- |
| $\alpha$ | Learning rate | 0.012* | 7 in A&B (2011) |
| $\Gamma$ | Response threshold | 0.313* | 14 in A&B (2011) |
| $\rho$ | Input scaling factor | 1.764* | 7 (12 in A&B, 2011) |
| $\phi$ | Control signal scaling factor | 2.246* | 8 (13 in A&B, 2011) |
| $\psi$ | Mutual inhibition scaling factor | 0.724* | 8 (13 in A&B, 2011) |
| $\beta$ | Rate coding scaling factor | 0.0415 | 11 in A&B (2011) |
| $\sigma$ | Variance of noise in control units | 0.0013 | 11 in A&B (2011) |
| $\theta$ | Risk aversion parameter | 1.0* | 1 in B&A (2017) |
| $B$ | Baseline level of excitation in $\omega$ units | 1.0 | 1, 2 |
| $K$ | Input threshold to the control units | 0 | 7, 8 |
| $W^C_{slope}$ | Slope of the stimuli-to-control weights | 0.3 | |
| $W^C_{intercept}$ | Intercept of the stimuli-to-control weights | -0.5 | |
| $W^F_{i-to-i}$ | Representation-to-control weights for the same option | -1.3 | |
| $W^F_{i-to-j}$ | Representation-to-control weights for the other option | 0.2 | |

**Figure S2.**
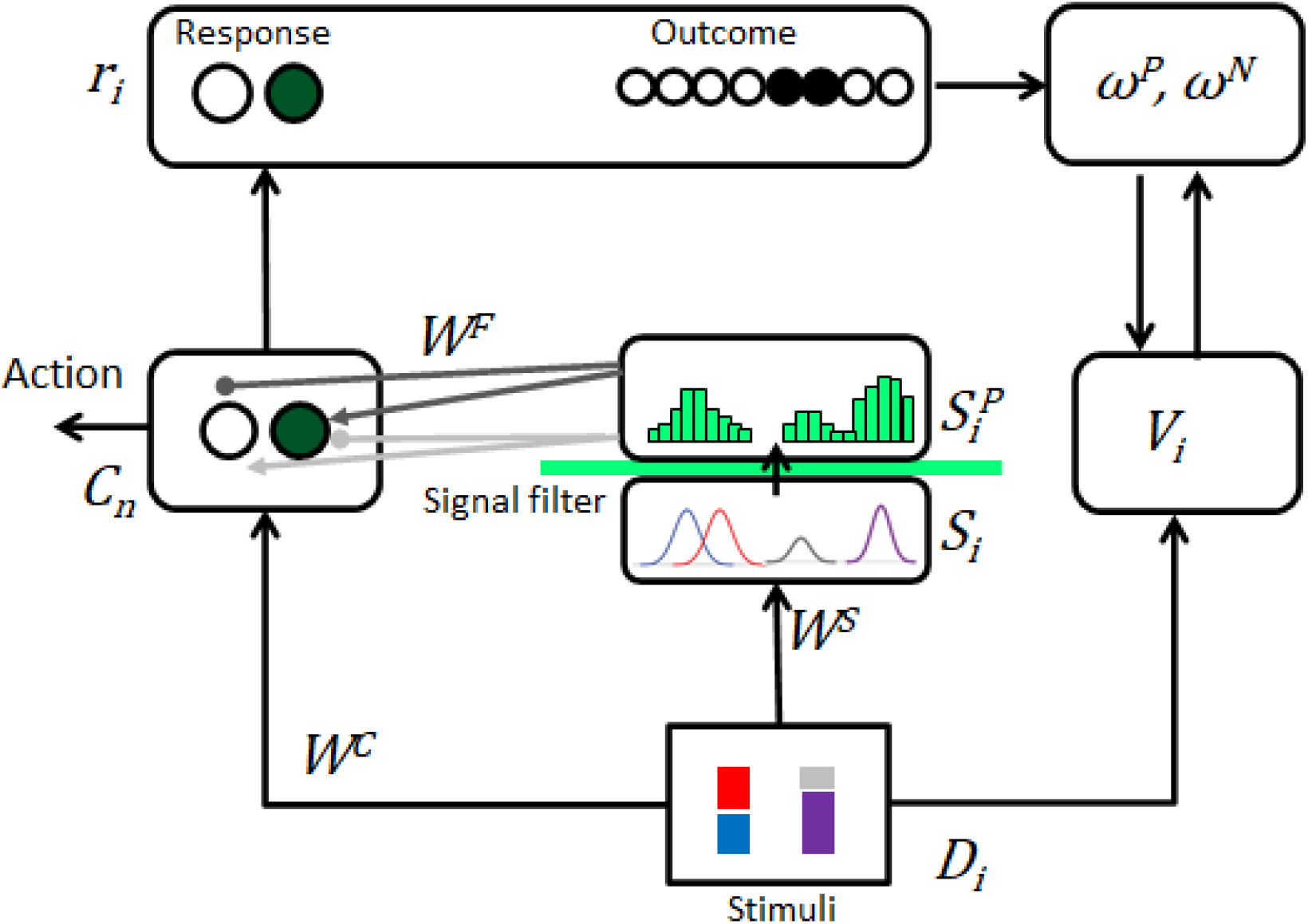
Monkey: PRO model as configured to accept inputs and generate responses for the monkey behavioral task.

**Table S2.** Model Parameters (Human)

| Parameter | Description | Value | Equation |
| --- | --- | --- | --- |
| $\alpha$ | Learning rate | 0.012* | 7 in A&B (2011) |
| $\Gamma$ | Response threshold | 0.313* | 14 in A&B (2011) |
| $\rho$ | Input scaling factor | 1.764* | 7 (12 in A&B, 2011) |
| $\phi$ | Control signal scaling factor | 2.246* | 8 (13 in A&B, 2011) |
| $\psi$ | Mutual inhibition scaling factor | 0.724* | 8 (13 in A&B, 2011) |
| $\beta$ | Rate coding scaling factor | 1.038* | 11 in A&B (2011) |
| $\sigma$ | Variance of noise in control units | 0.005* | 11 in A&B (2011) |
| $\theta$ | Risk aversion parameter | 1.0* | 1 in B&A (2017) |
| $B$ | Baseline level of excitation in $\omega$ units | 1.0 | 1, 2 |
| $K$ | Input threshold to the control units | 1.79 | 7, 8 |
| $W_{Bound}^F$ | A constant bound to the representation-to-control weights $W^F$ | 0.30 | 9 |
| $Y_{norm, 1}$ | Valence normalization parameter when tokens are won | 100 | 11 |
| $Y_{norm, 2}$ | Valence normalization parameter when tokens are lost | 36 | 11 |

**Figure S2.**
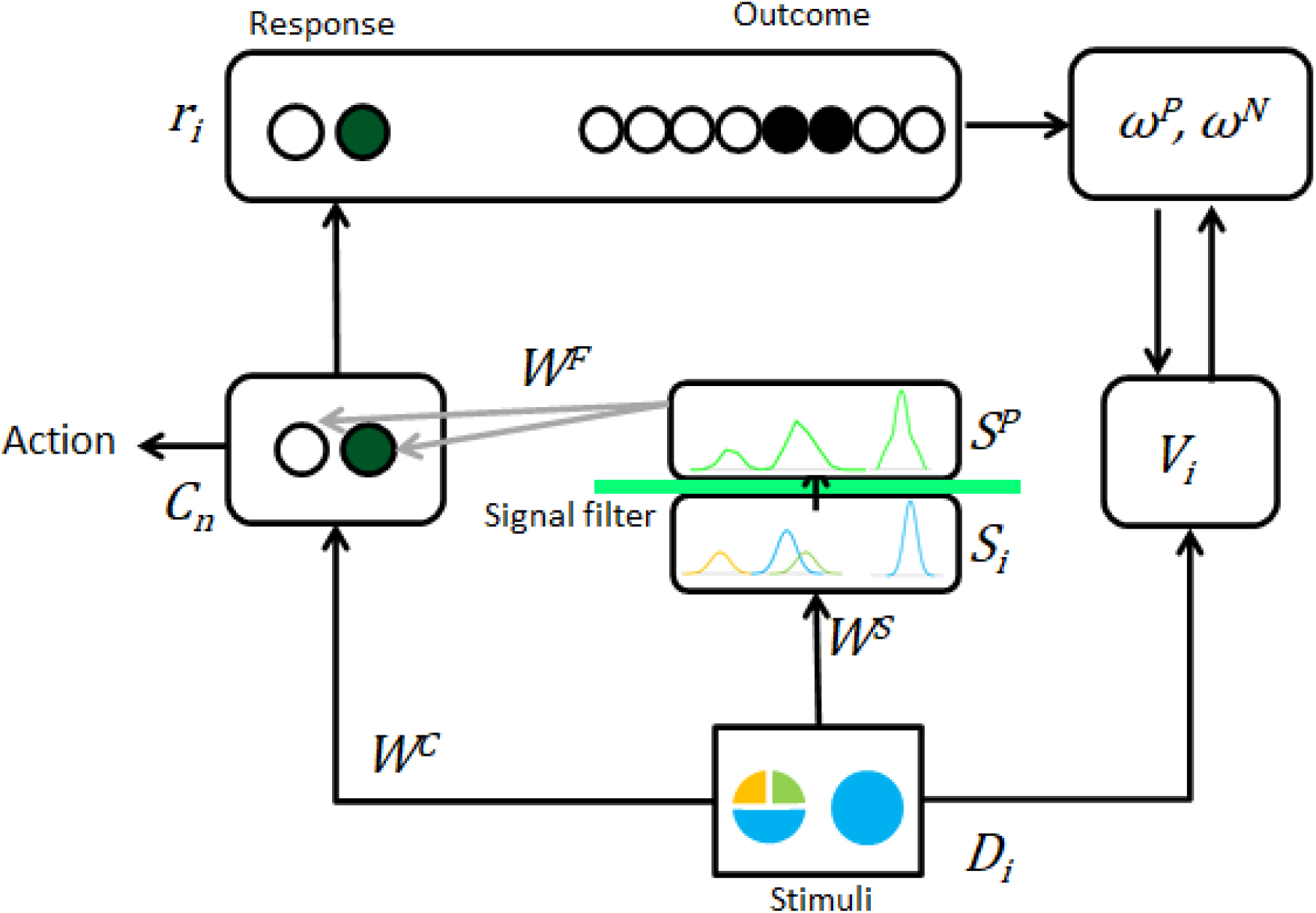
Human: PRO model as configured to accept inputs and generate responses for the human behavioral task.

#### 4. Open source simulation code

The modified PRO control model is available on GitHub: https://github.com/CogControlLab/PROControl

## References

Alexander, W. H., & Brown, J. W. (2011). Medial prefrontal cortex as an action-outcome predictor. Nature Neuroscience, 14(10). https://doi.org/10.1038/nn.2921

Alexander, W. H., & Brown, J. W. (2014). A general role for medial prefrontal cortex in event prediction. Frontiers in Computational Neuroscience, 8. https://doi.org/10.3389/fncom.2014.00069

Alexander, W. H., & Brown, J. W. (2019). The Role of the Anterior Cingulate Cortex in Prediction Error and Signaling Surprise. Topics in Cognitive Science, 11(1). https://doi.org/10.1111/tops.12307

Amador, N., Schlag-Rey, M., & Schlag, J. (2000). Reward-Predicting and Reward-Detecting Neuronal Activity in the Primate Supplementary Eye Field. Journal of Neurophysiology, 84(4). https://doi.org/10.1152/jn.2000.84.4.2166

Azab, H., & Hayden, B. Y. (2017). Correlates of decisional dynamics in the dorsal anterior cingulate cortex. PLOS Biology, 75(11). https://doi.org/10.1371/journal.pbio.2003091

Azab, H., & Hayden, B. Y. (2018). Correlates of economic decisions in the dorsal and subgenual anterior cingulate cortices. European Journal of Neuroscience, 47(8), 979–993.

Azab, H., & Hayden, B. Y. (2020). Partial integration of the components of value in anterior cingulate cortex. Behavioral Neuroscience, 134(4), 296.

Blanchard, T. C., & Hayden, B. Y. (2014). Neurons in dorsal anterior cingulate cortex signal postdecisional variables in a foraging task. Journal of Neuroscience, 34(2), 646–655.

Blanchard, T. C., Strait, C. E., & Hayden, B. Y. (2015). Ramping ensemble activity in dorsal anterior cingulate neurons during persistent commitment to a decision. Journal of Neurophysiology, 114(4), 2439–2449.

Bliss-Moreau, E., Santistevan, A. C., Bennett, J., Moadab, G., & Amaral, D. G. (2021). Anterior cingulate cortex ablation disrupts affective vigor and vigilance. Journal of Neuroscience, 41(38), 8075–8087.

Boroujeni, K. B., Watson, M., & Womelsdorf, T. (2021). Gains and Losses affect Learning Differentially at Low and High Attentional Load. BioRxiv, 2009–2020.

Botvinick, M. M., Braver, T. S., Barch, D. M., Carter, C. S., & Cohen, J. D. (2001). Conflict monitoring and cognitive control. Psychological Review, 108(3). https://doi.org/10.1037/0033-295X.108.3.624

Braver, T. S., Gray, J. R., & Burgess, G. C. (2012). Explaining the Many Varieties of Working Memory Variation: Dual Mechanisms of Cognitive Control. In Variation in Working Memory. https://doi.org/10.1093/acprof:oso/9780195168648.003.0004

Brown, J. W., & Alexander, W. H. (2017). Foraging Value, Risk Avoidance, and Multiple Control Signals: How the Anterior Cingulate Cortex Controls Value-based Decisionmaking. Journal of Cognitive Neuroscience, 29(10). https://doi.org/10.1162/jocn_a_01140

Brown, J. W., Reynolds, J. R., & Braver, T. S. (2007). A computational model of fractionated conflict-control mechanisms in task-switching. Cognitive Psychology, 55(1), 37–85.

Carter, C. S. (1998). Anterior Cingulate Cortex, Error Detection, and the Online Monitoring of Performance. Science, 280(5364). https://doi.org/10.1126/science.280.5364.747

Ebitz, R. B., & Hayden, B. Y. (2016). Dorsal anterior cingulate: a Rorschach test for cognitive neuroscience. Nature Neuroscience, 19(10), 1278–1279.

Farashahi, S., Donahue, C. H., Hayden, B. Y., Lee, D., & Soltani, A. (2019). Flexible combination of reward information across primates. Nature Human Behaviour, 3(11), 1215–1224.

Fukunaga, R., Purcell, J. R., & Brown, J. W. (2018). Discriminating Formal Representations of Risk in Anterior Cingulate Cortex and Inferior Frontal Gyrus. Frontiers in Neuroscience, 12. https://doi.org/10.3389/fnins.2018.00553

Gemba, H., Sasaki, K., & Brooks, V. B. (1986). “Error” potentials in limbic cortex (anterior cingulate area 24) of monkeys during motor learning. Neuroscience Letters, 70(2), 223–227. https://doi.org/10.1016/0304-3940(86)90467-2

Heilbronner, S., & Hayden, B. (2013). Contextual factors explain risk-seeking preferences in rhesus monkeys. Frontiers in Neuroscience, 7, 7.

Heilbronner, S. R., & Hayden, B. Y. (2016). Dorsal anterior cingulate cortex: a bottom-up view. Annual Review of Neuroscience, 39, 149–170.

Hayden, B. Y., Heilbronner, S. R., Pearson, J. M., & Platt, M. L. (2011). Surprise Signals in Anterior Cingulate Cortex: Neuronal Encoding of Unsigned Reward Prediction Errors Driving Adjustment in Behavior. Journal of Neuroscience, 31(11). https://doi.org/10.1523/JNEUROSCI.4652-10.2011

Holroyd, C. B., & Yeung, N. (2012). Motivation of extended behaviors by anterior cingulate cortex. In Trends in Cognitive Sciences (Vol. 16, Issue 2, pp. 122–128). Elsevier Current Trends. https://doi.org/10.1016/j.tics.2011.12.008

Jahn, A., Nee, D. E., Alexander, W. H., & Brown, J. W. (2016). Distinct Regions within Medial Prefrontal Cortex Process Pain and Cognition. The Journal of Neuroscience, 36(49). https://doi.org/10.1523/JNEUROSCI.2180-16.2016

Kennerley, S. W., Behrens, T. E. J., & Wallis, J. D. (2011). Double dissociation of value computations in orbitofrontal and anterior cingulate neurons. Nature Neuroscience, 14(12).https://doi.org/10.1038/nn.2961

Kolling, N., Behrens, T. E. J., Mars, R. B., & Rushworth, M. F. S. (2012). Neural Mechanisms of Foraging. Science, 336(6077). https://doi.org/10.1126/science.1216930

Maisson, D. J.-N., Cash-Padgett, T. V, Wang, M. Z., Hayden, B. Y., Heilbronner, S. R., & Zimmermann, J. (2021). Choice-relevant information transformation along a ventrodorsal axis in the medial prefrontal cortex. Nature Communications, 12(1), 1–14.

Markowitz, H. (1952). Portfolio Selection. The Journal of Finance, 7(1), 77–91. https://doi.org/10.2307/2975974

Matsumoto, M., Matsumoto, K., Abe, H., & Tanaka, K. (2007). Medial prefrontal cell activity signaling prediction errors of action values. Nature Neuroscience, 10(5). https://doi.org/10.1038/nn1890

Medalla, M., & Barbas, H. (2009). Synapses with Inhibitory Neurons Differentiate Anterior Cingulate from Dorsolateral Prefrontal Pathways Associated with Cognitive Control. Neuron, 61(4), 609–620. https://doi.org/10.1016/j.neuron.2009.01.006

Nieder, A. (2002). Representation of the Quantity of Visual Items in the Primate Prefrontal Cortex. Science, 297(5587). https://doi.org/10.1126/science.1072493

Niki, H., & Watanabe, M. (1979). Prefrontal and cingulate unit activity during timing behavior in the monkey. Brain Research, 171(2), 213–224. https://doi.org/10.1016/0006-8993(79)90328-7

Parvizi, J., Rangarajan, V., Shirer, W. R., Desai, N., & Greicius, M. D. (2013). The will to persevere induced by electrical stimulation of the human cingulate gyrus. Neuron, 80(6), 1359–1367. https://doi.org/10.1016/j.neuron.2013.10.057

Paus, T. (2001). Primate anterior cingulate cortex: where motor control, drive and cognition interface. Nature Reviews Neuroscience, 2(6), 417–424.

Shenhav, A., Botvinick, M. M., & Cohen, J. D. (2013). The expected value of control: An integrative theory of anterior cingulate cortex function. In Neuron (Vol. 79, Issue 2, pp. 217–240). Cell Press. https://doi.org/10.1016/j.neuron.2013.07.007

Strait, C. E., Blanchard, T. C., & Hayden, B. Y. (2014). Reward value comparison via mutual inhibition in ventromedial prefrontal cortex. Neuron, 82(6), 1357–1366.

Strait, C. E., Sleezer, B. J., Blanchard, T. C., Azab, H., Castagno, M. D., & Hayden, B. Y. (2016). Neuronal selectivity for spatial positions of offers and choices in five reward regions. Journal of Neurophysiology, 115(3), 1098–1111.

Walton, M. E., Bannerman, D. M., Alterescu, K., & Rushworth, M. F. S. (2003). Functional Specialization within Medial Frontal Cortex of the Anterior Cingulate for Evaluating Effort-Related Decisions. The Journal of Neuroscience, 23(16). https://doi.org/10.1523/JNEUROSCI.23-16-06475.2003

Zarr, N., & Brown, J. W. (2016). Hierarchical error representation in medial prefrontal cortex. Neuroimage, 124, 238–247.

## References

Brown, J. W., & Alexander, W. H. (2017). Foraging Value, Risk Avoidance, and Multiple Control Signals: How the Anterior Cingulate Cortex Controls Value-based Decision-making. Journal of Cognitive Neuroscience, 29(10). https://doi.org/10.1162/jocn_a_01140

